# The unknown lipids project: harmonized methods improve compound identification and data reproducibility in an inter-laboratory untargeted lipidomics study

**DOI:** 10.1101/2023.02.01.526566

**Authors:** Tong Shen, Ciara Conway, Kaitlin R. Rempfert, Jennifer E. Kyle, Sean M. Colby, David A. Gaul, Hani Habra, Fanzhou Kong, Kent J. Bloodsworth, Douglas Allen, Bradley S. Evans, Xiuxia Du, Facundo M. Fernandez, Thomas O. Metz, Oliver Fiehn, Charles R. Evans

## Abstract

Untargeted lipidomics allows analysis of a broader range of lipids than targeted methods and permits discovery of unknown compounds. Previous ring trials have evaluated the reproducibility of targeted lipidomics methods, but inter-laboratory comparison of compound identification and unknown feature detection in untargeted lipidomics has not been attempted. To address this gap, five laboratories analyzed a set of mammalian tissue and biofluid reference samples using both their own untargeted lipidomics procedures and a common chromatographic and data analysis method. While both methods yielded informative data, the common method improved chromatographic reproducibility and resulted in detection of more shared features between labs. Spectral search against the LipidBlast in silico library enabled identification of over 2,000 unique lipids. Further examination of LC-MS/MS and ion mobility data, aided by hybrid search and spectral networking analysis, revealed spectral and chromatographic patterns useful for classification of unknown features, a subset of which were highly reproducible between labs. Overall, our method offers enhanced compound identification performance compared to targeted lipidomics, demonstrates the potential of harmonized methods to improve inter-site reproducibility for quantitation and feature alignment, and can serve as a reference to aid future annotation of untargeted lipidomics data.

## Introduction

Liquid chromatography-mass spectrometry-based lipidomics is the current method of choice for comprehensive analysis of the lipidome in biological samples.^1^ Through its development over the past two decades, lipidomics has been strengthened by large-scale efforts such as LIPID MAPS^2,3^ and the Lipidomics Standard Initiative,^4^ which have helped establish consistent nomenclature, online databases, data analysis tools and best practices for lipidomics research. Lipidomics can be performed using untargeted or targeted approaches, which use high resolution-accurate mass instruments or tandem or hybrid quadrupole mass spectrometers, respectively. Targeted lipidomics is often implemented using commercial kits provisioned with internal standards and pre-configured methods.^5^ Several ring-trial studies have demonstrated good reproducibility of these methods across laboratories.^6,7^ The primary tradeoff of targeted lipidomics is the limitation of the analytes to those pre-configured in the method, which prevents detection of new or unpredicted compounds and the discovery of unexpected biological insights that might result from such data.

Untargeted lipidomics uses high-resolution MS with data-dependent or data-independent tandem mass spectrometry and unbiased feature extraction to detect and characterize both expected and unknown lipids. Untargeted lipidomics methods typically allow detection of more lipids than targeted methods in a single analysis, and sensitivity can be comparable to targeted approaches, especially those that attempt to assay hundreds of lipids or more.^8^ However, few efforts have been made to develop standardized techniques for untargeted lipidomics and associated data processing. As a result, it is difficult to aggregate untargeted lipidomics data from multiple sources to validate trends or perform meta-analyses. Furthermore, unknown features are prevalent in untargeted lipidomics data and lipid identification remains a major bottleneck. Modern spectral libraries such as LipidBlast^9^ and software packages such as NIST MS Search^10^, NIST MSPepSearch^11^, or MS-DIAL^12^ can help identify lipids using MS/MS spectra,^9^ but even with the best available spectral search tools, more lipid features typically remain unidentified than identified. In addition to novel lipids of biological interest, unknown features may include many artifacts such as in-source fragments, adducts, contaminants, and non-lipid molecules. This diversity complicates data interpretation, especially since no satisfactory effort has been made to systematically catalog both known and unknown lipids observed in specific sample types and across different laboratories.

Here, we describe an untargeted lipidomics ring trial focused on assessing the extent of overlap of both identified and unknown lipids in data from five expert laboratories. The study workflow, illustrated in Fig 1, began with multi-site LC-MS/MS analysis of four pooled human plasma reference samples provided by NIST and one sample each of human skeletal muscle, bovine heart and liver, and porcine brain. All samples were prepared for analysis using a common protocol including use of 72 isotopically labeled internal standards. Each site analyzed extracts using their own standard LC-MS lipidomics protocol, henceforth termed “in-house LC-MS data acquisition”, and also performed “common method LC-MS data acquisition” that proscribed use of a specific column and gradient but allowed use of any high resolution-accurate mass instrument. Data analysis, including feature detection, alignment and annotation using MS/MS database search was first performed both separately by each lab, termed “in-house data processing”. Data from all labs were also processed together in a single MS-DIAL project to generate a master alignment table, which was termed the“common data processing pipeline”. Ion mobility spectrometry-mass spectrometry data were acquired at a single site and aligned with the LC-MS/MS data from all labs. The primary outcome of the study was a comparison of inter-site compound identification consistency and unknown detection, yielding insight into the current state of inter-site reproducibility in untargeted metabolomics. Our results also highlight the potential of harmonized chromatographic methods and common data analysis pipelines to improve reproducibility, enhance compound identification performance, and to help derive more biological insight from untargeted lipidomics data.

**Fig. 1:**
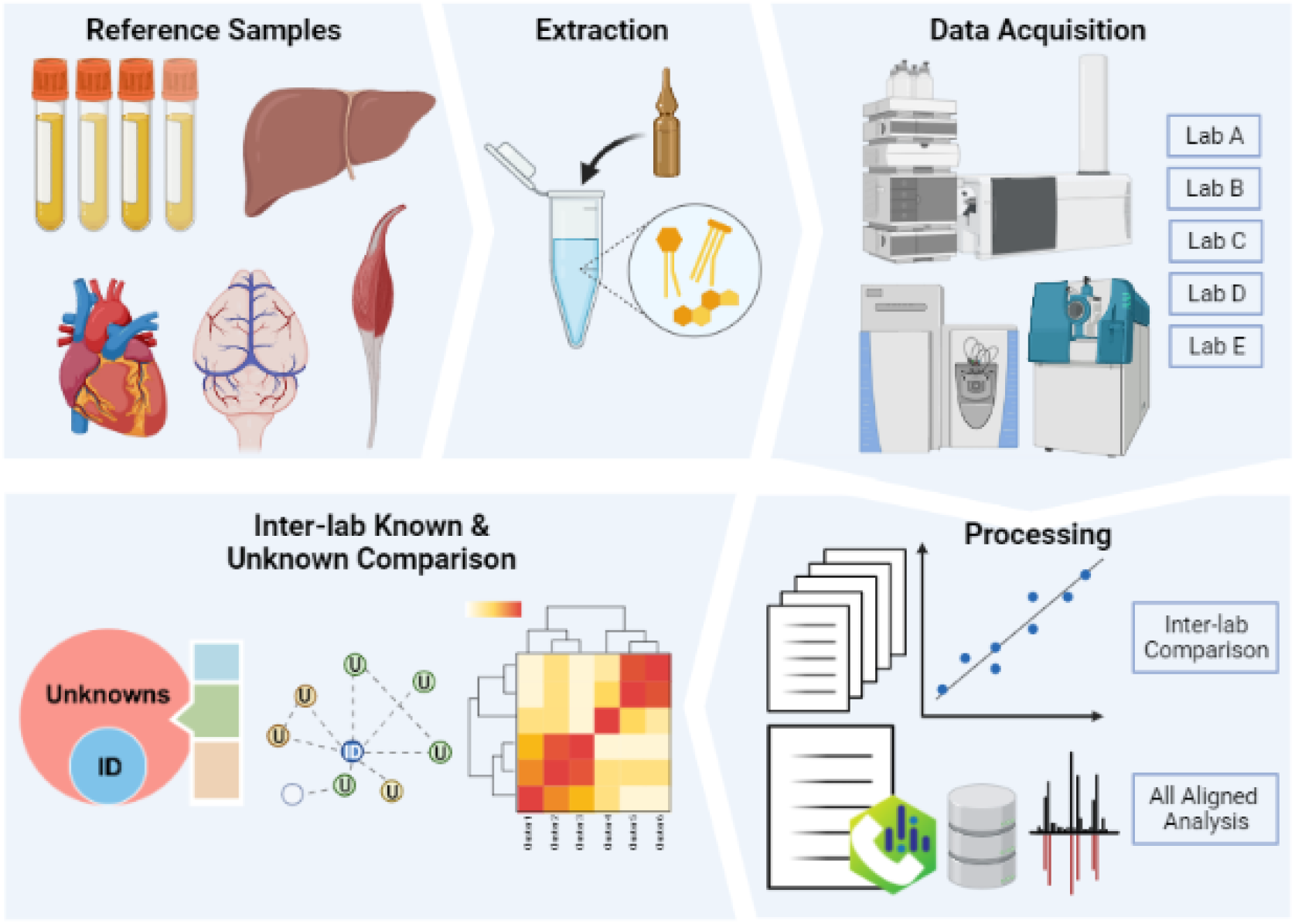
Inter-laboratory untargeted lipidomics study workflow

## Results

### Inter-site retention time reproducibility

To evaluate chromatographic reproducibility between study sites, measured retention time (RT) values for the 72 lipid internal standards were compared. RT data from Lab A, which used the common method exclusively, were compared to in-house methods (Fig. 2A). Lipid elution order was largely consistent between labs, but due to the use of distinct columns and gradient profiles, RT correlations were non-linear. Second-order polynomial regression yielded curve fits with R^2^ values of 0.98 or higher. When the common method was used (Fig. 2B), an excellent linear correlation was observed, with R^2^ values for a least-squares linear fit exceeding 0.997 for all datasets (Fig. 2A). Lab C’s measured RT correlation showed slightly divergent slope; this was attributed to use of a quaternary gradient UHPLC pump with low-pressure solvent mixing, which had a larger delay volume than the high-pressure mixing binary pumps used by other labs. Measured RTs of internal standards were used to adjust retention times of all other peaks in the dataset, resulting in RT standard deviation of < 0.1 min across all labs.

**Fig. 2:**
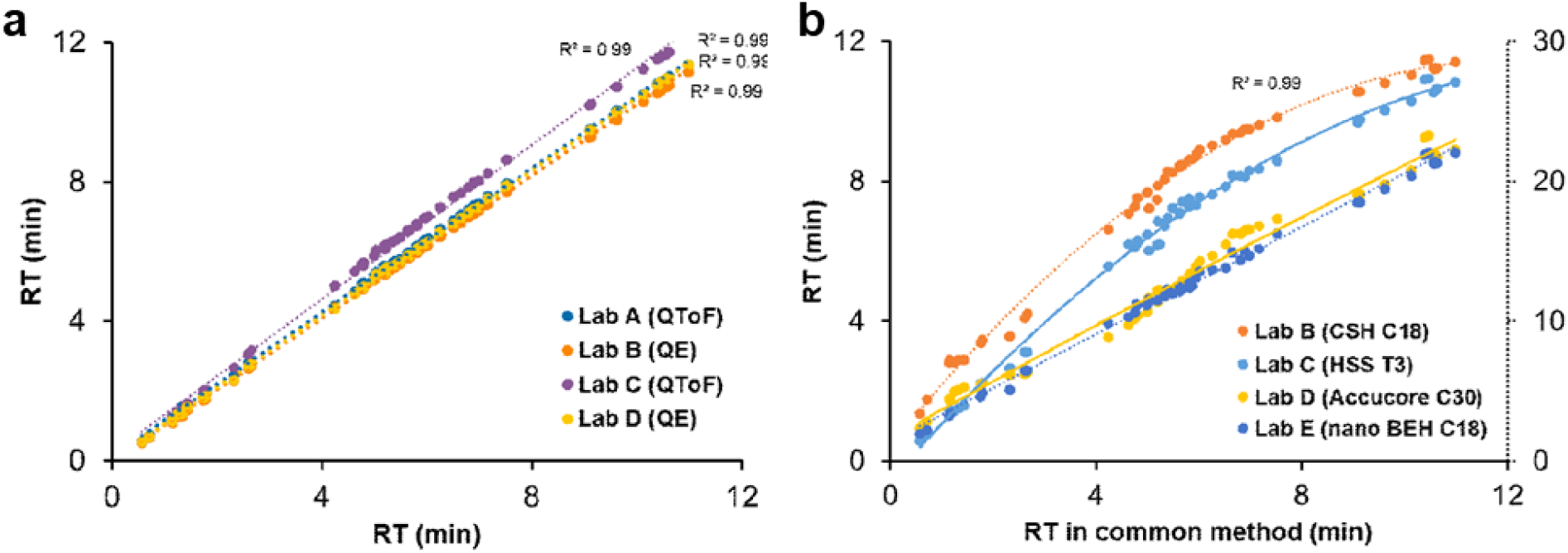
Inter-laboratory retention time alignment. Retention time correlation for 72 stable isotope-labeled lipid standards between five laboratories using **a** in-house-method data acquisition and **b** common-method data acquisition

### Comparison of in-house versus common-pipeline lipidomics data processing

Next, untargeted feature detection and alignment were performed as described in the methods. The data from in-house LC-MS acquisition methods were processed only by the lab that acquired the data, whereas the common-method LC-MS data were processed both separately by each site (in-house data processing) and using the common-pipeline method. The common-pipeline data analysis required accommodations compared to typical data processing parameters. First, minimum peak height for feature detection was set to a low value, 1E4, that permitted detection of low-abundance features in Q-TOF data, resulting in a lower-than typical threshold for the Orbitrap data. Second, gap filling was turned off in MS-DIAL processing to avoid artificial reporting of abundance values for features initially detected by only one site or instrument type. These parameters resulted in higher prevalence of missing values compared to a conventional, single-lab data processing workflow. Thus, for most cross-site data comparison of the data, a missingness filter was applied to remove features not detected in at least 50% of the samples in one or more tissue types. This had the effect of requiring a feature be detected by at least three labs, as each site ran equal replicates of each sample.

Table 1 lists summary statistics for feature detection using both the common-pipeline and the in-house data processing strategies. The four separate feature tables generated by individual labs using in-house processing were systematically aligned by m/z, RT and relative abundance using Metabcombiner,^13^ which generated a master feature table comparable to the common-pipeline processing performed in MS-DIAL, which are included with the manuscript as Supplementary Data 1. The total number of features detected using each processing method ranged from ∼20,000 to ∼65,000. Common-pipeline processing resulted in a greater proportion of features detected and aligned by at least two labs than in-house data processing (6404 vs 5350 for positive ion and 6121 vs 1909 for negative ion). Likewise, for identified lipids, the common-pipeline method aligned roughly twice as many features across labs in both positive mode and negative mode. Using the common-pipeline data processing strategy, 42% of features detected at the MS1 level were assigned an MS2 spectrum in positive ion mode and 37% in negative mode.

**Table 1.** Summary of detected and aligned features, identified lipids, and unknown features in **a** common-pipeline data and **b** in-house data

|  | Common-pipeline data processing (MS-DIAL) |  |
| --- | --- | --- |
|  | (+) ion | (-) ion |
| Total feature count | 27,454 | 65,381 |
| Feature count, <50% missingness in 1+ tissues | 6404 | 6121 |
| Total identified lipids | 1951 | 1140 |
| Identified lipids <50% missingness | 1572 | 952 |
| Unknowns w/ MS2, <50% missingness | 2043 | 1890 |
| Unknowns w/o MS2, <50% missingness | 2789 | 3279 |

|  |  | In-house data processing + external feature alignment (MetabCombiner) |  |
| --- | --- | --- | --- |
|  |  | (+) ion | (-) ion |
| Total feature count |  | 36,844 | 22,241 |
| Features aligned between labs: | 4 labs | 794 | 292 |
|  | 3 labs | 1560 | 508 |
|  | 2 labs | 2996 | 1,129 |
|  | 1 lab (unaligned) | 31494 | 20,312 |
| Total identified lipids |  | 2794 | 846 |
| Identified lipids **aligned** between labs: | 4 labs | 135 | 120 |
|  | 3+ labs | 222 | 162 |
|  | 2+ labs | 395 | 219 |
|  | 1 lab (unaligned) | 2078 | 345 |

Feature detection and alignment using the in-house LC-MS/MS data, for which each lab used its own processing software and protocol, was substantially more variable. Total feature counts for each site ranged from approximately 2000 to over 20,000 per ion mode (Supplementary Data 2). The number of features which could be confidently aligned between labs was lower, with 85% and 91% of features observed by only one lab in positive and negative ion modes, respectively.

### Degenerate feature annotation

Dgenerate features including in-source fragments and adducts are widely observed in untargeted lipidomics data, but their prevalence has not been compared between labs. For initial annotation of degenerate features, the common LC-MS method / common data analysis pipeline data were processed using Binner.^14^ This Java software tool groups features into bins by retention time, annotates probable adducts and in-source fragments, including additive combinations of degeneracies, based on a user-supplied “mass shift” table. It also calculates inter-feature correlation of coeluting species to allow detection of unpredicted degeneracies. Probable degenerate features assigned by Binner were flagged in the master feature list (Supplemental Data 1) but not removed, as cataloging of consistently-observed adducts and in-source fragments in lipidomics data was one of the aims of the study. Of the 6404 features with 50% overall missingness or less, 606 (∼9%) features in positive ion mode and 1219 (∼20%) were annotated as probable degenerate features based on the mass shift annotation. An additional 612 features in positive ion mode and 659 (11%) in negative ion mode had a high correlation (Pearson’s R >0.9) with a co-eluting feature. Of predicted adducts, approximately two-thirds (403/606) were observed in four or all five of the datasets in the common method/common pipeline alignment table, suggesting that degeneracy formation is a relatively consistent phenomenon across laboratories, at least for features abundant enough to be consistently detected.

As an additional strategy to detect in-source fragments, we mined the unknowns with MS2 data using ISFinder as described in the methods to detect co-eluting features with similar patterns of fragmentation. Using this strategy, 360 additional putative in-source fragments were detected in positive mode and 63 in negative mode in the common dataset. Overall, the total proportion of features annotated as degenerate was 25% for positive ion mode and 32% for negative mode. The proportion of degenerate features is lower than reported some comparable lipidomics studies^15^ in spite of a relatively rigorous degeneracy annotation approach. One possible explanation is that our inter-laboratory data alignment may act as a pre-filter, resulting in a set of features with a greater proportion of unique compounds and mostly consistently-observed degenerate features.

### Identified lipid sample composition

Using compound identification criteria described in the methods, the common-pipeline data analysis identified 1951 unique lipids representing 43 lipid classes in positive mode and 952 unique lipids representing 59 lipid classes in negative ion mode. Lipid names and classes were assigned according to the LIPID MAPS Lipid Classification System^16^ and the ClassyFire taxonomy schema.^17^ The resulting class distribution of identified lipids, illustrated in Figs. 3B and 3E, shows modest variability between sample types. Brain tissue contained a higher proportion of sphingolipids, whereas sterol lipids, one of the least frequently detected lipid classes, were more numerous in plasma. The identified lipid composition can be further refined to the lipid subclass level as illustrated in Fig 3C and 3F. We also used unsupervised principal component analysis (PCA) to assess qualitative differences in lipid abundance between groups, illustrated in Fig. S1 A clear separation was observed between different tissue types; plasma samples clustered closely together but the distinct pooled samples could still be distinguished. A substantial separation was observed between data acquired using QTOF and Orbitrap instruments, implying that relative signal differences between instrument types remain detectable in the data following normalization by log transformation and autoscaling. Visualization of the data as a heatmap with hierarchical clustering by sample type (Figure S2) revealed additional sample-specific patterns in lipid abundance.

**Fig. 3:**
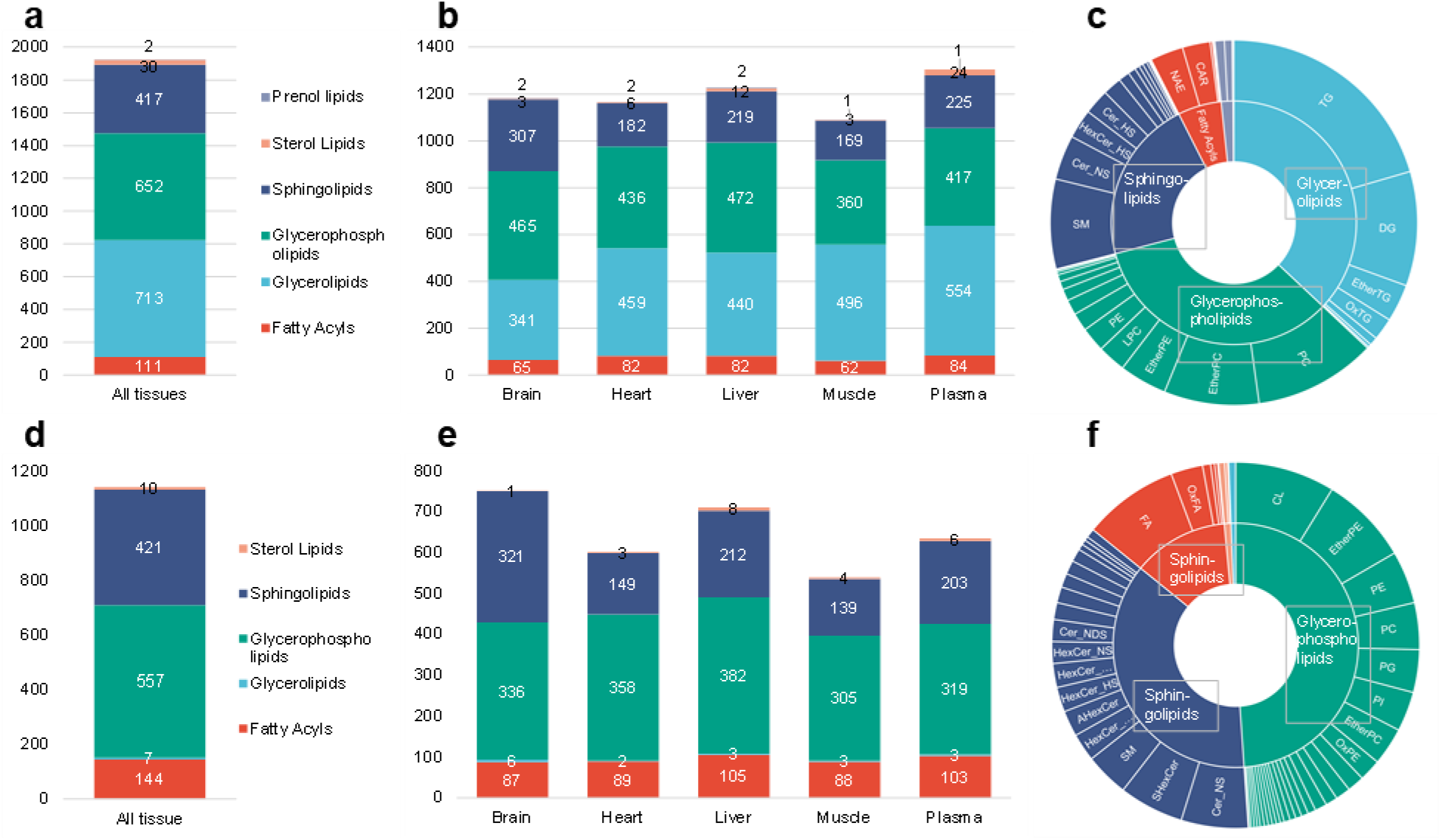
Identified lipid class distribution. **a** positive ion mode superclass distribution for identified lipids for the complete dataset, **b** superclass distribution sorted by tissue type **c** lipid class level distribution for all tissues, **d-f** same as above for negative ion mode data

### Chromatographic and spectral properties of identified lipids and unknown features

To visually evaluate the spectral and chromatographic behavior of identified and unknown lipids, RT vs. m/z plots were created as illustrated in Fig. 4A. As previously reported,^12^ lipids group by class when visualized in this manner. In certain regions of the plot many abundant lipid classes overlap (PC, PE, CE), in others regions only one lipid species is detected (e.g. CL and FAFHA), and in other regions no identified lipid species are found. When unknown features are added (Fig. 4B), some fall in the same regions as the identified lipids while others appear in regions where no major identified lipids are observed, suggesting that these unknowns may represent contaminants, background signal, or non-lipid species. A comparable analysis was performed using ion mobility-mass spectrometry (IMMS) data, which was used to generate collision cross section (CCS) vs. m/z plots (Fig. 4C). Identified lipids exhibited a distinctive CCS vs. m/z correlation, with different lipid classes (e.g. glycerophospholipids and glycerolipids) occupying unique space within this series of trendlines; as in the RT vs m/z plot, unknown features occupied a wider range of CCS vs. m/z conformational space than known features. These observations suggest a strategy for prioritizing follow-up analysis of unknowns that “conform” to chromatographic and spectral properties of previously identified lipids. To support predictive analysis of unknowns, we fit a two-parameter power law regression model to the observed CCS vs. m/z trend of assigned lipids and generated a 99% predictive interval (PI) for the model, representing the CCS range expected for 99% of measured lipids at a given m/z. Of the 6119 unknown features with measured CCS values, 3551(58%) fell within the 99% PI and were flagged as tentative lipid-like structures in (**Supplementary Data 1**).

**Fig. 4:**
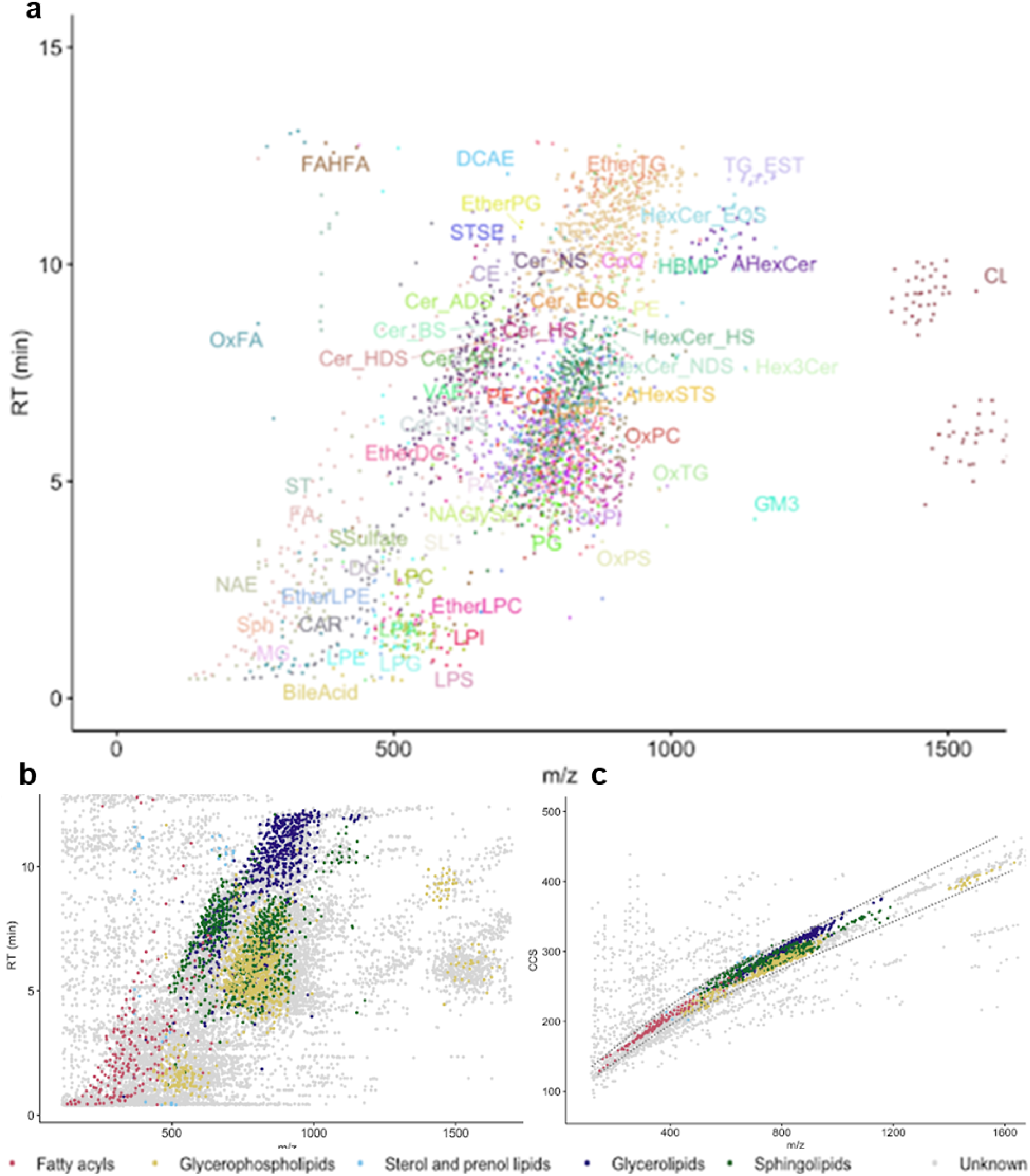
Chromatographic and Spectral Distribution of Identified and Unknown Lipids. **a** RT vs m/z distribution of identified lipids, colored by lipid class **b** RT vs. m/z distribution of identified (colored by superclass) and unidentified (gray) features. **c** Comparison of CCS vs m/z distribution of identified (colored by superclass) and unknown (gray) features with the 99% prediction interval for identified lipids plotted as dotted lines. Features shown in all plots are those detected in >50% of samples of at least one tissue type.

We also assessed the overlap of identified lipids and unknown features between sample types, illustrated as Venn diagrams in Fig. 5. For identified lipids, the largest single grouping was for lipids found in all five tissue types (327 and 182 for positive and negative ion mode, respectively). In contrast, the largest groupings of unknown features were those observed in only one sample type, including 1057 features observed only in muscle in positive ion mode and 1018 observed only in brain in negative ion mode. When overlapping features were compared within the four distinct reference human plasma samples, a higher proportion of both identified and unknown features were detected in all or most of the plasma samples, although features detected in only one sample type were also observed (Fig. S3).

**Fig. 5:**
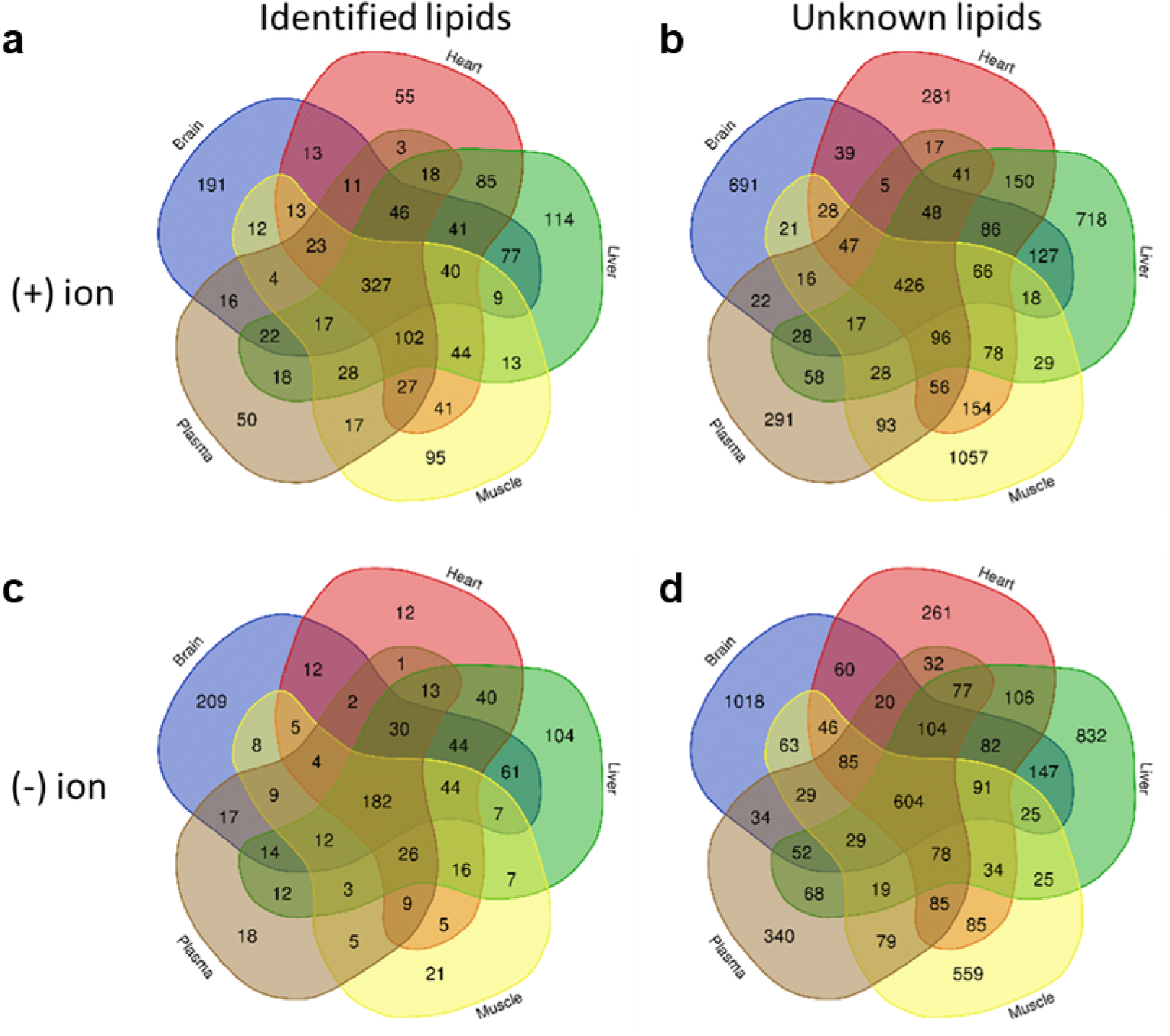
Overlap of identified and unknown lipids by tissue type. Data shown for features detected in >50% of common-method runs (including data from all study sites) of at least one tissue type.

### Automated classification of unknown features

To classify the unknown features and predict potential structural motifs of the compounds they represent, we used an automated strategy which combined NIST identity and hybrid searching performed via NIST MSPepSearch (http://chemdata.nist.gov)^18^ and in-silico fragmentation via MS-FINDER^19,20^ as described in the methods; a visual schema is illustrated in Fig. 6A-B. The automated spectral classification schema successfully assigned compound class to the majority of unidentified features with MS2 spectra. At the compound superclass level (Fig. 6C), the largest single set of unknown features were classified as lipids and lipid-like molecules, though less than half of the total set of unknown features fell within this group. Other well-represented superclasses included organic nitrogen/oxygen-containing compounds, benzenoids, and organoheterocyclic compounds. Among those compounds classified as lipids the most frequently observed lipid classes were glycerophospholipids, glycerolipids and fatty acyls; (Fig. 6D). Notably, features classified as sphingolipids were substantially enriched in brain tissue compared to all other tissue types, consistent with findings from the known lipid data.

**Fig. 6:**
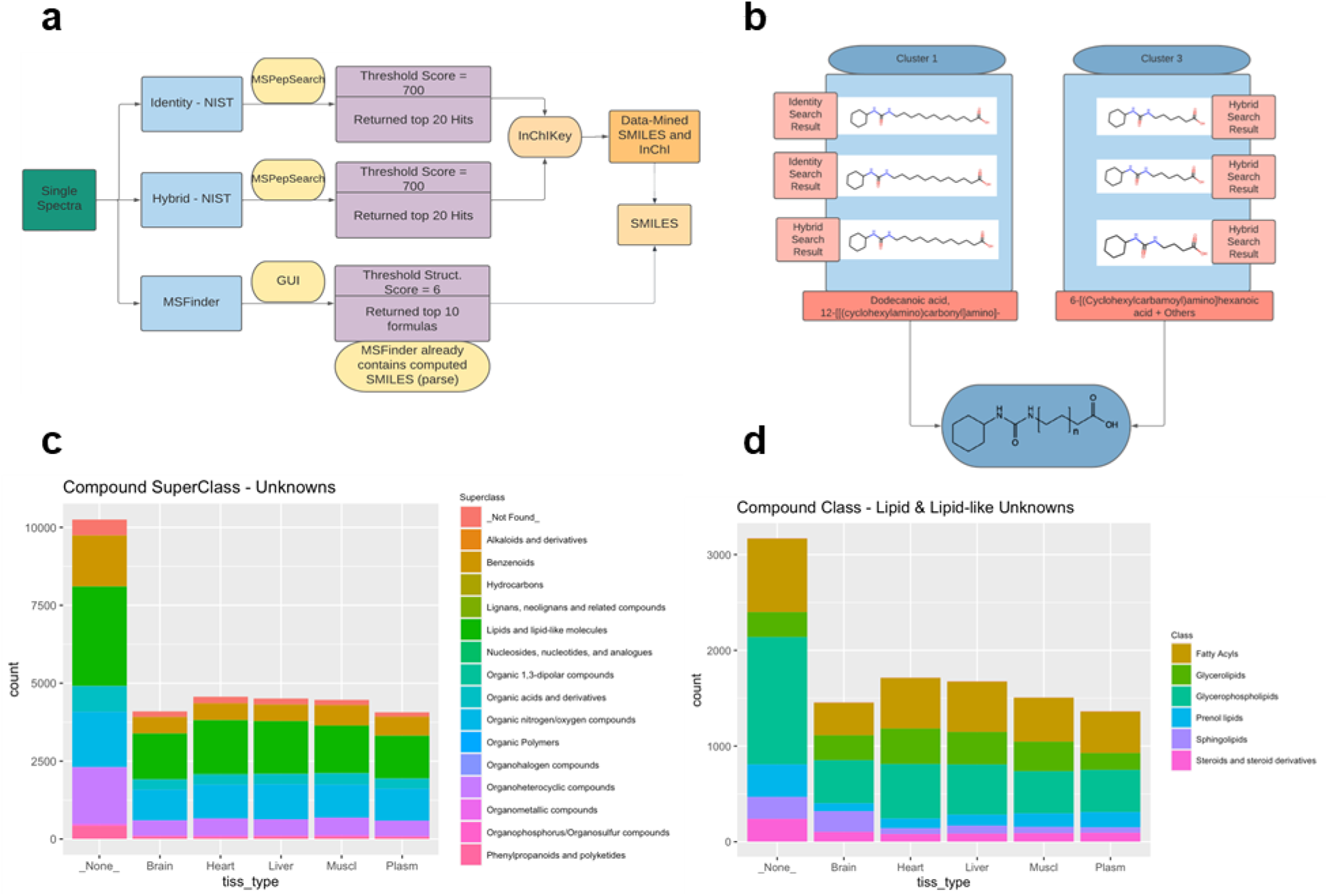
Unknown feature classification using spectral search. **a** Schema for classification of unknown features using multiple search strategies. **b** Representative example of structural prediction for an unknown lipid. **c** Unknown feature class distribution. **d** Lipid superclass distribution for subset of lipid-like unknowns

### Spectral networking analysis

As an additional approach to characterize unknown features, spectral networking analysis was performed on MS2 spectra from the study using the Global Natural Products Social Molecular Networking database (https://gnps.ucsd.edu). Processing resulted in the generation of 192 distinct molecular networks with feature counts ranging from 94 to 2 features per network. As illustrated in Fig. 7, spectral network families contained both identified and unknown features, but were often dominated by metabolites within a single class or several closely related classes such as triglycerides (Fig. 7A) or sphingomyelins and hexosylceramides (Fig. 7B). Features linked by edges were observed to differ by acyl chain length and desaturation as well as other chemical modifications such as oxidation. Triglycerides were widely distributed through all tissue types, but other spectral networks were dominated by species found in a single tissue. Hexosylceramides are glycosylated lipids with recently recognized critical roles in various essential cellular functions,^21^ which in our study were mostly observed in brain tissue. Spectral networks also highlighted exceptions, such as the lipid identified as Hex2Cer(18:1/16:0), a dihexosylceramide that was observed in muscle and plasma.

**Fig. 7:**
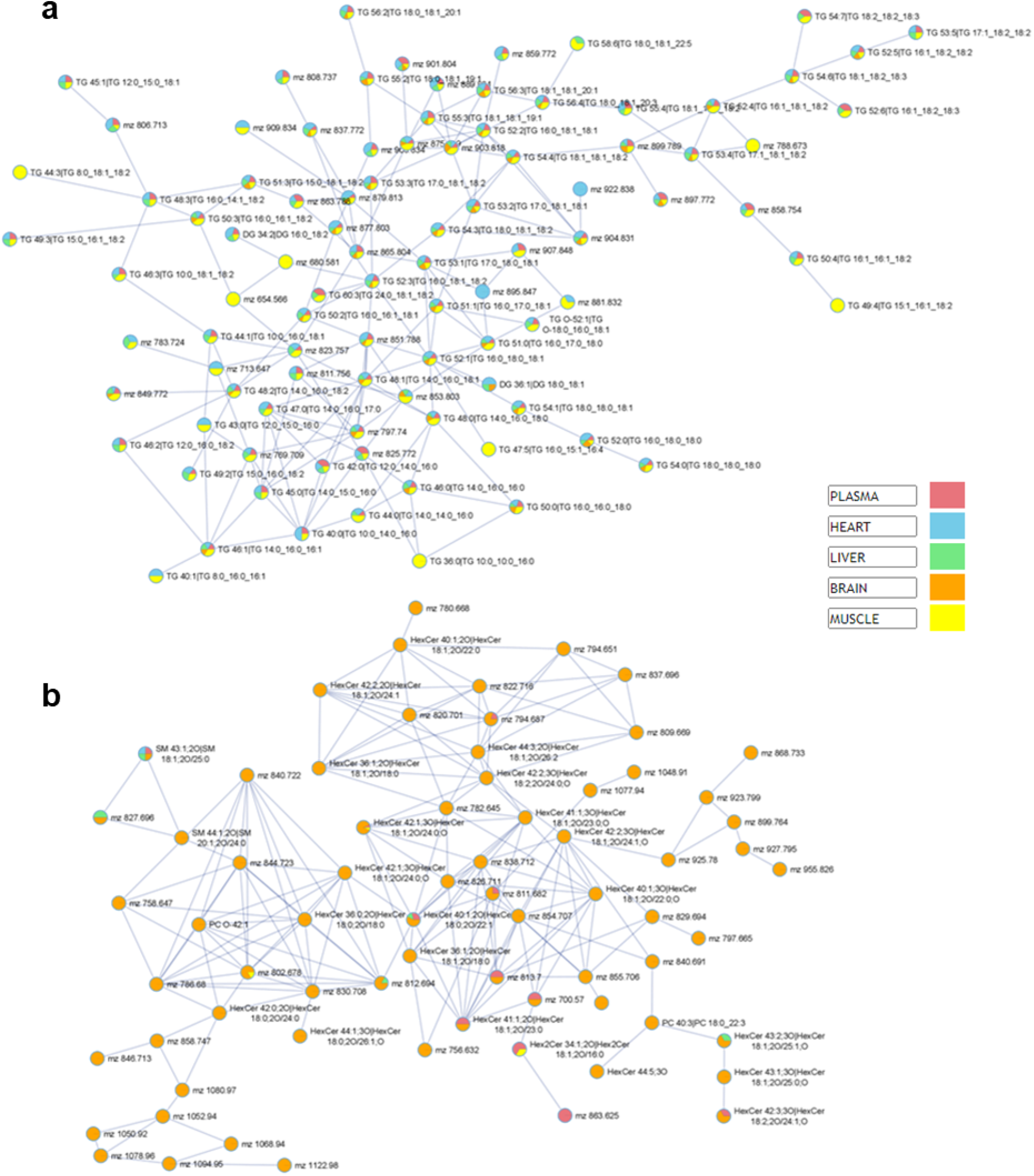
Representative spectral family networks generated using molecular networking analysis in GNPS. ^43^ **a** triglyceride-associated network observed in multiple tissues **b** hexosylceramide network observed primarily in brain tissue

### Identification of uncommon and novel lipid species

Some annotated lipids and unknowns were observed across all or most sample types, whereas others were tissue-specific, as expected. The porcine brain showed the highest number of unique annotated lipids, particularly hexosylceramides, as noted above. Another rare class of lipids detected in our data were sulfonolipids. This unique class of sphingolipids features sulfonic acid group in the sphingoid base and has previously been reported in the gastrointestinal microbial lipidome.^22^ Among the samples we investigated, we observed the highest abundance of sulfonolipids in bovine liver, which to our knowledge represents the first report of such species in mammalian tissue using an untargeted lipidomics approach. Another unique class of lipids annotated in our data were oxidized phospholipids and TGs, which showed distinct differential abundance between different matrices. Detected oxygenated lipids ranged from 1O to 3O, and represented mostly hydroperoxide- and hydroxide-modifications. OxPE were highest in abundance among the annotated oxidized lipids. The pooled plasma from hypertriglyceridemia subjects had very high levels of TG species, as expected, but surprisingly was not rich in oxTG. OxTG and unmodified TG showed differential levels among the samples, which is indicative of physiological oxidation rather than autoxidation. Such observations were consistent among all the labs.

By integrating results from our NIST identity search, NIST Hybrid Search, MS-FINDER, and GNPS spectral networking strategies, we were able to predict the compound superclass and class of most of the detected unknown features with MS/MS spectra. Many of the unknowns were, as expected, classified as lipid and lipid like molecules. Among this category, the main predicted lipid classes are glycerophospholipids followed by fatty acyls. The next most common compound classes were organic nitrogen and oxygen-containing compounds, organic heterocyclic compounds, then prenol lipids, steroids, and steroid derivatives. The latter classes are intriguing, as such hits were very limited among the annotated lipids. Such classes are scarcely reported or annotated in untargeted lipidomics data due to their limited presence in spectral libraries, especially rule-based *in silico* libraries such as LipidBlast. In spite of their smaller size, libraries based on authentic standards such as NIST20 are complementary to lipid in-silico libraries, especially for compound classes where in-silico spectral prediction is not as effective. Although chemical structures of the hybrid search results could not be used to directly identify unknown species, in some cases, these results could be extended to assign a definitive sub-structure, and in other instances the search results they generated helped shed light on the presence of unpredicted modified lipids. Manual review of hybrid search results helped lead to identification of oxygen-modified lipids that are not included in in-silico libraries, *e*.*g*. PC 34:2 (1O) and FA 20:4 (O3).

As reported previously, a portion of the unknowns represent degenerate features. Adduct ions are not limited to MS1 spectra but can also be observed as fragment ions within MS/MS spectra. Such adduct-specific fragmentation patterns are not fully incorporated into rule-based MS/MS libraries such as Lipidblast. PCs, LPCs and either PCs were easily annotated as protonated ion forms in positive mode, yet their sodiated adducts were initially characterized as unknown features. The lack of adduct-incorporating fragmentation rules led to low score or even no hits for spectral similar-scoring near-isobaric compounds (*e*.*g*. LPC O-16:1 vs. LPC P-16:0 [M+Na]+). The hybrid search also revealed unknowns that were likely to be in-source fragments of annotated lipids. For example, a long chain base structure which eluted at the retention time range of phospholipids and sphingolipids appeared to be a sphingosine in-source fragment of several ceramides, in accordance with ISFinder results.

## Discussion

Untargeted lipidomics depends on accurate compound identification to enable biologically valid interpretation of the data. Historically, compound identification methods and reporting criteria have varied from lab to lab, resulting in inconsistent data quality. Recent community-led initiatives have called for more rigorous and consistent criteria for lipid identification and reporting^23^ and proposed a minimum reporting checklist.^24^ In this study, we compared lipidomics data generated using typical lab-selected protocols with data generated using specific shared chromatographic and data analysis methods. Using both strategies, we analyzed a muti-tissue set of standard reference materials and investigated to what extent different labs running different instruments identify the same lipids and observe the same (or different) unknown features. Our findings highlight several areas in which inter-laboratory harmonization can increase data quality and reduce investigator effort for untargeted lipidomics studies.

Our untargeted data alignment approach allowed an unprecedented evaluation of unknown feature overlap between laboratories. We considered separately those unknowns for which a data-directed MS2 spectrum was collected and those for which it was not. For unknown features without an MS2 spectrum, the majority were not observed consistently (with <50% missingness in one or more tissue types) in more than one laboratory. These features are mostly low-abundance, and often represent contaminants or artifacts. For those which are not, identification is likely to require targeted MS2 acquisition or other strategies to improve signal. Contrastingly, more than 55% of unknowns with an MS2 spectrum were observed by multiple laboratories (2530/4592 and 3539/6375 for positive and negative ion modes, respectively}. Features with MS2 spectra that were not observed by multiple labs may represent background peaks and contaminants unique to a specific lab, or degeneracies (adducts and in-source fragments) unique to a specific instrument or ion source. Notably, the inter-laboratory study design served as an effective data-filtering strategy. Unknowns observed consistently by multiple labs can be prioritized for identification, as they are more likely to represent a true unidentified lipid.

An innovative feature of our approach was use of a common chromatographic method accompanied by a multi-class set of isotopically labeled lipids for retention time correction. Feature alignment within a dataset is typically achieved by retention time warping or binning methods implemented in software such as XCMS,^25,26^ MZmine^27^ or MS-DIAL^12,28^. However these approaches are less well-suited to aligning data acquired over multiple months or by distinct laboratories, which suffer from less consistent feature detection and larger shifts in retention and relative intensity. While software tools have been developed to assist with inter-batch ^29–32^ and inter-lab^13^ feature alignment, the use of a fixed chromatographic method with a set of readily-detectable internal standards spread throughout the entire chromatogram offers superior performance. In our study, this approach gave post-correction retention time differences <0.1 min for all compounds, compared to post-correction alignment ranging between 0.1-0.5 min for lab-selected protocols. Our strategy could easily be adapted to generate retention indices rather than adjusted retention times, allowing inclusion of reference retention information into spectral databases.

Another novel aspect of our study was the common data processing pipeline, in which raw data files from different labs were co-batched and aligned in MS-DIAL. Comparing only data collected using the common chromatographic method, the common-pipeline data analysis approach aligned and identified over twice as many lipids between two or more labs in both positive and negative ion modes (Table 1). More unknown features were also aligned using the common-pipeline analysis approach. Our data generated among the highest total number of identified lipids in an untargeted study to date, and enabled identification of unknowns from multiple classes ranging from common triglycerides to unique oxidized lipids and sulfonlyated species.

The data from this study may also serve to help validate lipid identifications or prioritize unknown features in future lipidomics datasets, particularly if the common chromatographic method is used for sample analysis. While it was not the intent of this study to produce a formal database of lipid features, the success of our inter-laboratory feature alignment demonstrates the feasibility of tissue-specific, experimentally-derived feature databases containing both known lipids and unknown features, which would include retention times and/or retention indices generated using an established, reproducible chromatographic method. Such a resource would greatly facilitate compound identification and reduce duplication of effort currently associated with annotating unknown features in untargeted lipidomics data.

By leveraging harmonized instrumental analysis and data processing methods, our study demonstrates that untargeted lipidomics can achieve identification accuracy and inter-laboratory consistency comparable to targeted methods with superior lipid compound class coverage, while retaining the ability to discover and identify unknown lipids. Further efforts are required to evaluate the performance of our common-method approach with larger, clinical-scale lipidomics studies and meta-studies. With these improvements, and with sufficient community agreement, we expect lipid annotation accuracy can be increased and effort reduced. Standardized methods, common internal standards, and common reference materials can be expected to play a major role in such efforts.

## Methods

### Samples and Materials

Bovine heart and liver total lipid extract, porcine brain total lipid extract and UltimateSPLASH™ ONE internal standard mixture were purchased from Avanti Polar Lipids (Birmingham, AL). Additional internal standard compounds were purchased from Sigma-Aldrich (St. Louis, MO). Extraction solvents, mobile phase additives were all LC-MS grade and were purchased by individual participating labs from multiple chemical vendors.Plasma samples were candidate human plasma reference materials (RM 8231 – Frozen Human Plasma Suite) obtained from the National Institute of Standards and Technology (NIST), which included three pooled plasma samples representing different subject populations and health states (hypertriglyceridemia pool, diabetic pool, and young African American pool). For this study, these samples were designated as P1, P2, and P3, respectively. NIST SRM 1950 human pooled plasma was also included as a common reference material widely used in the lipidomics and metabolomics community;^33^ this was designated sample P4. Human skeletal muscle extract was prepared from a bulk pool of frozen tissue obtained from de-identified consented human donors.

Samples were compiled at a single site, extracted, aliquoted, dried under nitrogen gas, and distributed to all participating labs via shipping on dry ice. Samples were maintained at -80C until the time of extraction and/or analysis.

### Sample preparation

Plasma samples were prepared by liquid-liquid extraction with methyl tertiary butyl ether (MTBE) using a modified version of the Matyash protocol.^34^ Twenty uL of plasma in a microcentrifuge tube were vortexed with 975 uL of ice-cold 3:10 methanol:MTBE containing a mix of 76 internal standards (Table S1). After 5 minutes on a shaker at 4°C, 188 uL water were added, samples were vortexed again for 20 sec and then centrifuged 2 min at 14,000 rcf. Three hundred and fifty μL of the upper (organic) phase were transferred to a clean tube and dried by centrivap and stored at -20°C until analysis. Samples were resuspended in 110 uL of 9:1 methanol:toluene and transferred to autosampler vials for analysis. Human skeletal muscle was extracted using the same procedure, substituting 20 mg of frozen tissue for the plasma and reconstituting in 200uL of 9:1 methanol:toluene. For tissue total lipid extracts purchased from Avanti Polar Lipids (heart, liver, brain), 10 uL of extract was transferred to an autosampler vial, spiked with internal standards (Table S1), dried under nitrogen gas and stored at -80C for distribution and analysis. Dried samples were reconstituted in 200 uL 9:1 methanol:toluene on the day of analysis.

### LC-MS/MS

Samples were analyzed two ways by participating labs: using a common method and using in-house methods. For the **common method LC-MS/MS analysis**, labs selected their own high-resolution accurate-mass LC-MS/MS instruments and mass spectrometer settings (detailed in Supplementary Information) but used a common chromatographic method. The column was a Waters CSH (2.1 mm x 100 mm; 1.7 μm particle size) with a matched guard column. The column was maintained at 65ºC and a flow rate of 600 μl/min. For positive ion mode runs, mobile phase A was (60:40) acetonitrile:water with 10 mM ammonium formate and 0.1% formic acid; mobile phase B was (10:90) acetonitrile:isopropanol with 10 mM ammonium formate and 0.1% formic acid. For negative ion mode runs, mobile phase A was acetonitrile:water (60:40) with 10 mM ammonium acetate; mobile phase B was (10:90) acetonitrile:isopropanol with 10 mM ammonium acetate. The gradient profile for both ion modes was as follows (min, %B): 0,15%; 2,30%; 2.5,48%; 11,82%; 11.5,99%; 12,99%; 12.1,15%. Total run time was 16 minutes. Individual labs selected their own mass spectrometer and its operating parameters. For the **in-house method LC-MS/MS analysis**, each lab selected its own chromatographic method, mass spectrometer, and operating parameters, which are described in Supplementary Information.

### Data Analysis and compound identification

Data analysis was performed using two pipelines: via a common data processing pipeline and using in-house methods. For the **common pipeline data analysis**, raw data from all participating labs were converted to a universal file format using Reifycs ABFconveter, then loaded together as a single project (for each ion mode) into the data analysis tool MS-DIAL 4.70.^12,28^. MS-DIAL was used to perform feature detection and alignment and was configured with the following analysis parameters: Data collection MS1 tolerance 0.01 Da, MS2 tolerance 0.025Da; retention time correction enabled (programmed with m/z and RT of spiked internal standards, Table S1); minimum peak height 1000, linear moving average peak smoothing enabled at level 3, minimum peak width 9 scans; sigma window value 0.5; MS/MS abundance cut off 0; adduct joining disabled, alignment RT tolerance 0.1 min, alignment MS1 tolerance 0.015 Da, gap filling disabled. MS-DIAL selects the MS/MS spectrum to represent a feature from the data file which produced the highest-scoring match. If no match was produced, the spectrum from the precursor with highest abundance was selected. For compound identification, the Lipidblast database distributed with MS-DIAL was used with a score threshold of 700 and precursor mass accuracy of < 0.01Da.

The resulting aligned feature table contained m/z values, retention times, MS/MS data (if available), peak height, and compound annotations. When specific acyl chain configurations were reported, identifications were manually reviewed to ensure spectral support existed for these assignments.

For **in-house data analysis**, feature detection, alignment, quantitation, and compound identification were performed according to the standard operating procedures of the labs in which data were acquired. Required criteria included that all identifications be made either using MS/MS data or an accurate mass and RT match with an authentic standard analyzed in-house using the same method, and that the data be reported in a single aligned feature table containing at minimum: m/z, retention time, identification or annotation if available, and relative quantitation by peak height. Details regarding each lab’s in-house data analysis methods are provided in the Supplementary Information. The software tool MetabCombiner^13^ was used to align features by m/z, RT and relative abundance between the in-house datasets generated by different labs. Operating parameters and detailed methods used for Metabcombiner are in Supplementary Information.

### Identification of in-source fragmentation by ISFinder

One of the caveats of mass-spectrometry-based lipidomic studies is the artifacts generated from in-source fragmentation (ISF) . Failure to recognize such artifacts often leads to false compound identification and erroneous biological interpretation.^35^ Therefore, a novel automatic detection method, ISFinder, was implemented to explicitly identify ISFs. The workflow is an improvement upon the method by Guo et al.^36^, with modifications including the incorporation of spectral entropy similarity scoring to improve accuracy and robustness of fragment identification.^37^ ISfinder is programmed in Python 3.8 and freely available on GitHub (https://github.com/lancelot0821/ISFinder). For analysis using ISFinder, spectra were cleaned by removing both the precursor peak and low-signal peaks (less than 0.5% relative intensity), followed by normalizing feature abundance to the sum of all peak intensities. Non-informative spectra with a calculated spectral entropy less than 0.5 were also removed.^37^ The ISFinder algorithm is built upon the following premises: 1) ISFs should coelute with their corresponding precursors; 2) ISFs are stable and abundant; and 3) ISFs and their precursors are essentially the same chemical with similar fragmentation patterns.^36^ For each unknown, all identified compounds within 3 seconds of its retention time are flagged as precursor candidates. Further filtering is applied to require the presence of unknown feature’s m/z in the MS2 spectrum of the precursor candidates. Before calculating the entropy similarities, the MS2 spectra of precursor candidates are truncated so that only the ions below the unknown’s m/z are considered. An entropy similarity score of 0.6 or higher between the unknown and truncated precursor spectra was empirically determined to be a suitable threshold above which the unknown should be considered an ISF of the candidate precursor. The performance of ISFinder was validated on compounds with known fragmentation by loss of water, demonstrating an accuracy of 94%.

### Statistical analysis and plotting

For multivariate statistics, data were loaded into Metaboanalyst.^38,39^ Features with >50% missing values were removed and missing values were replaced by estimated limit of detection (⅕ of the minimum positive value for each feature). Data were log-transformed and autoscaled (mean-centered and divided by the standard deviation of each feature). Plots were prepared using Microsoft Excel and R Statistical Software v4.2. ^40^ Principal component analysis was performed using the prcomp R package.

### Unknown feature classification using MSPepSearch and MS-FINDER

Unidentified features represented by m/z and RT in tabular format (Supplementary Data 1) were searched using NIST MSPepSearch^41^ in identity search mode against a compiled “.MSP” file containing all MS2 spectra from the study to align each unknown compound with its MS2 spectra using RT and m/z with tolerances of 0.1 min and 0.008 Da, respectively. A new .MSP file containing all the unknown spectra was passed into both MSPepSearch and MS-FINDER. The similarity score threshold cutoff for MSPepSearch was set to 700 for both identity and hybrid, while the MS-FINDER similarity threshold cutoff was set to 6.0. Illustrated in Fig 5A, this workflow generated a list of up to 20 candidate hits for each search approach per unknown compound. These results were piped into a structural similarity comparison workflow in which similar compounds were grouped through hierarchical clustering with complete linkage, using the Tanimoto Similarity Score as the distance metric.^42^ The similarity cutoff was set at 85% for inclusion in the same cluster of compounds. Each result was required to have an associated SMILE that was converted to the 2D fingerprint notation. The results from MSPepSearch contained only the InChIKey and not the corresponding SMILE string. Thus, for these entries, the InChIKey was searched against the PubChem Database, through a REST API, to obtain the matching SMILE. For MS-Finder, the SMILE string was automatically generated. Within the workflow, each compound’s results were clustered and then loaded into ClassyFire in order to obtain the taxonomy of each result, both the top3 and top1 clusters were analyzed to determine if a general structural classification could be deduced. All hits are included in the Supporting Information, Table S2.

### Spectral Networking Analysis

Two spectral networks, one for positive ion mode and one for negative, were created using the molecular networking workflow on the Global Natural Products Social Molecular Networking (GNPS) server, (http://gnps.ucsd.edu).^43^ Since GNPS generates networks based on similarity between experimental spectra, data files from only the single lab with the largest MS/MS dataset (Lab C) were used as an input to ensure consistent spectral features and fragmentation patterns. The data was filtered by removing all fragment ions within +/-17 Da of the precursor m/z. MS/MS spectra were also window-filtered by choosing only the top 6 fragment ions in the +/-50Da window throughout the spectrum. The precursor ion mass tolerance and fragment ion tolerance was set at 0.02 Da. The minimum cosine score for edges in the network was set at 0.7 and 4 or more matched peaks were required. Edges between two nodes were kept in the network only if each of the nodes appeared in each other’s respective top 10 most similar nodes. The maximum size of a molecular family was set to 100, and the lowest scoring edges were removed from molecular families until the molecular family size was below this threshold. Spectra in the network were searched against GNPS public spectral libraries using a minimum cosine score of 0.5 and 4 or more matched peaks. The resulting data were exported as a Cytoscape network, and features in the Cytoscape network were re-aligned with the master feature list (Supplemental Data 1) using MetabCombiner as described previously. Feature names from the aligned data were imported into Cytoscape and used as the default feature labels for network visualization; GNPS-assigned feature names were used if no MS-DIAL identification was available.

## Code Availability

Code for ISFinder software is available on GitHub (https://github.com/lancelot0821/ISFinder).

## Acknowledgements

We gratefully acknowledge the National Institute of Standards and Testing for provision of the human plasma candidate reference material, and Tracey Schock and Christina Jones for discussions and review of this manuscript. We acknowledge the NIH Common Funds Metabolomics Consortium and its participants for input into this work. Funding was provided by NIH Common funds grants ES030164, ES030158, ES030170, ES030167, and CA325507.

