## Supplementary Information for "The unknown lipids project: harmonized methods improve compound identification and data reproducibility in an inter-laboratory untargeted lipidomics study"

for

**Supplemental Methods**

**Lab A: LC-MS/MS conditions (Lab A), common method.** Lab A used only the **common LC-MS/MS method** for all sample analysis but implemented the method on two different mass spectrometers. Lab A instrument 1 was a ThermoFisher Q-Exactive Eactive Orbitrap mass spectrometer equipped with a Vanquish chromatograph. For analysis the electrospray ionization source was operated at a spray voltage of 3.6 kV for positive ionization mode and 3.0 kV for negative ionization mode, sheath, auxiliary, and sweep gas flows of 60, 25, and 2 (arbitrary units), respectively, and capillary temperature of 300°C and an aux gas heater temperature of 270°C. S-lens RF level was 50, AGC target was 1e6, and maximum inject time was 100ms. The instrument acquired full MS data with 120,000 resolution over a scan range of 120-1700 *m/z*. Lab A instrument 2 was an Agilent 6550 qTOF mass spectrometer equipped with an Agilent 1290 UPLC system. MS parameters were as follows: gas temp 325°C, drying gas flow 8L/min, nebulizer pressure 35 psig, sheath gas temp 350°C, sheath gas flow 11L/min. The capillary voltage was 3500V (for both positive and negative modes). MS acquisition was performed in full scan mode from m/z 60-1700Da with an acquisition rate of 2 spectra/sec, with reference mass correction enabled.

**Lab B: LC-MS/MS conditions, common method.** The total lipid extracts (TLEs) were analyzed by reversed-phase LC-ESI-MS/MS using a Waters Acquity UPLC H class system (Waters Corp., Milford, MA) coupled to a Q Exactive Orbitrap mass spectrometer (Thermo Scientific, San Jose, CA). The mass spectrometer was equipped with a Thermo HESI source. Its inlet was maintained at 300ºC while the HESI source was held at 370ºC with a spray voltage of 3.6kV for positive and 3.0kV for negative modes. The sheath, auxiliary, and sweep gas flows were 60, 25, and 2, respectively. A precursor scan of *m/z* 200-2000 at a mass resolution of 60k was followed by data-dependent MS/MS of the top 4 ions at a resolution of 15,000, a first fixed mass of *m/z* 70, and an isolation window of 1 *m/z*. Lipids were fragmented by HCD (higher-energy collision dissociation) using stepped normalized collision energies of 20, 30, and 40.

**Lab B: LC-MS/MS conditions, in-house method:** Samples were analyzed using the same instrument and mass spectrometer parameters as for the common method. The column used was a Waters CSH column (3.0 mm x 150 mm, 1.7 µm particle size) and separate over a 34-minute gradient (mobile phase A: ACN/H2O (40:60) containing 10 mM ammonium acetate; mobile phase B: ACN/IPA (10:90) containing 10 mM ammonium acetate) The gradient was as follows (time in min, %B): 0,40%; 2,50%; 3,60%; 12,70%; 15,75%; 17,78%; 19,85%; 22,92%; 25,99%; 34,99%; 34.5, 40%, 35,99%; 35.5,99%; 36,40%; 37,40%; 38, 40%. The flow rate was 0.25 mL/min except as follows (time in min, flow rate): 34.5,0.3 mL/min; 35,0.3 mL/min; 35.5,0.3 mL/min; 36,0.35 mL/min; 37,0.3 mL/min; 38,0.25 mL/min.

**Lab C: LC-MS/MS conditions, common method:** The total lipid extracts (TLEs) were analyzed by reversed-phase LC-ESI-MS/MS using an Agilent 1290 Infinitiy II quaternary pump (Agilent, Santa Clara, CA) coupled to an Agilent 6545 quadrupole time of flight mass spectrometer. LC parameters were as described in the main text. The mass spectrometer was equipped with an Agilent JetStream orthogonal electrospray ionization source maintained at the following parameters: nitrogen sheath gas flow, sheath gas temperature, drying gas flow, drying gas temperature, nebulizer pressure, and nozzle voltage at 10L\min, 275ºC, 5L\min, 325ºC, 30 psi, and 2kV respectively. . Its inlet was maintained at 300ºC while the HESI source was held at 370ºC with a spray voltage of 3.6kV for positive and 3.0kV for negative modes. The sheath, auxiliary, and sweep gas flows were 60, 25, and 2, respectively. A precursor scan of *m/z* 200-2000 at a mass resolution of 60k was followed by data-dependent MS/MS of the top 4 ions at a resolution of 15,000, a first fixed mass of *m/z* 70, and an isolation window of 1 *m/z*. Lipids were fragmented by HCD (higher-energy collision dissociation) using stepped normalized collision energies of 20, 30, and 40.

**Lab C: LC-MS/MS conditions, in-house method:** A Shimadzu CTO-20A Nexera X2 UHPLC systems equipped with a degasser, binary pump, thermostated autosampler, and a column oven was used for chromatographic separation. The column heater temperature was set at 55° C. 5 uL of lipid extract was injected onto a 1.8 µm particle diameter, 50 × 2.1 mm Acquity HSS UPLC T3 column (Waters, Milford, MA). Acetonitrile/water (40:60, v/v) with 10 mM ammonium acetate was mobile phase A and acetonitrile/water/isopropanol (10:5:85 v/v) with 10 mM ammonium acetate was mobile phase B. For chromatographic elution we used a linear gradient over a 20 min total run time, with 60% A and 40% B gradient in the first 10 minutes. Then the gradient was ramped in a linear fashion to 100% B which was maintained for 7 minutes. Thereafter the system was switched back to 60% B for 3 minutes. The flow rate was 0.4 mL/min The column was equilibrated for 3 min before the next injection. MS data acquisition was performed in both positive and negative ionization modes using a TripleTOF 5600 equipped with a Turbo VTM ion source (AB Sciex, Concord, Canada). The source voltage was set to 5500V for positive ionization and 4500V for negative ionization mode, the declustering potential was set to 60 V, and the source temperature to 450 C for both modes. The curtain gas flow, nebulizer, and heater gas was set to 30, 40, and 45 units respectively. The instrument performed one TOF MS survey scan (150 ms) and 15 MS/MS scans with a total duty cycle time of 2.4 s. The mass range in both modes was 50-1700 m/z. We controlled the acquisition of MS/MS spectra by data dependent acquisition (DDA) function of the Analyst TF software (AB Sciex, Concord, Canada) with the following parameters: dynamic background subtraction, charge monitoring to exclude multiply charged ions and isotopes, and dynamic exclusion of former target ions for 9 s. Rolling collision energy spread was set whereby the software calculated the collision energy value to be applied as a function of m/z. Mass accuracy was maintained by the use of an automated calibrant delivery system interfaced to the second inlet of the DuoSpray source.

**Lab D: LC-MS/MS conditions, common method:** LC-MS data were acquired using a Vanquish (ThermoFisher Scientific) chromatograph coupled to a high-resolution accurate mass Q-Exactive HF Orbitrap mass spectrometer (ThermoFisher Scientific). For analysis the electrospray ionization source was operated at a vaporizer temperature of 425°C, a spray voltage of 3.0 kV for positive ionization mode and 2.8 kV for negative ionization mode, sheath, auxiliary, and sweep gas flows of 60, 18, and 4 (arbitrary units), respectively, and capillary temperature of 275°C. The instrument acquired full MS data with 240,000 resolution over the 150-2000 *m/z* range. Samples were analyzed in random order with pooled QC injections collected at minimum every tenth injection. LC-MS/MS datawere acquired using a data dependent acquisition (DDA) strategy to aid in compound identification. $MS^2$ spectra were collected with a resolution of 120,000 and the dd-$MS^2$ were collected at a resolution of 30,000 and an isolation window of 0.4 *m/z* with a loop count of top 7. Stepped normalized collision energies of 10%, 30%, and 50% fragmented selected precursors in the collision cell. Dynamic exclusion was set at 7 seconds and ions with charges greater than 2 were omitted.

**Lab D: LC-MS/MS conditions, In-house method:** Instrumentation and MS parameters were identical to common method acquisition conditions. The chromatographic column used for in-house data acquisition was aThermoFisher Scientific Accucore™ C30 column (2.1 × 150 mm, 2.6 µm particle size),The mobile phases were 40:60 water:acetonitrile with 10 mM ammonium formate and 0.1% formic acid (mobile phase A) and 10:90 acetonitrile:isopropyl alcohol, with 10 mM ammonium formate and 0.1% formic acid (mobile phase B). The column temperature was set to 50°C, the injection volume was 2 µL, and the gradient program is as follows (min, %b) 0,20%; 1,60%; 5,70%; 8,90%, 8.2,100; 10.5,100; 10.7,20; 12,20.

**Lab C: LC-IMS-MS/MS conditions**. TLEs were analyzed using a Waters Acquity UPLC H class system (Waters Corp., Milford, MA) coupled with an Agilent 6560 Ion Mobility QTOF MS system (Agilent Technologies, Santa Clara, CA). LC parameters were identical to the common method described in the main text.  The Agilent 6560 used an Agilent JetStream orthogonal electrospray ionization source maintained at the following parameters: nitrogen sheath gas flow, sheath gas temperature, drying gas flow, drying gas temperature, nebulizer pressure, and nozzle voltage at 10L\min, 275ºC, 5L\min, 325ºC, 30 psi, and 2kV respectively. For the analysis, the IMS-MS inlet capillary was operated at 4kV, the high-pressure funnel maintained at 4.4 Torr, the trapping funnel at 3.8 Torr, and the rear funnel at 3.95 Torr. All funnels were operated at an RF of 100V DC. The IMS system was pressurized with ultrahigh purity nitrogen, and the drift potential was 1450V. MS/MS data were acquired during individual LC-IMS-MS/MS analyses at five fixed collision energies of 10, 20, 40, 60, and 80eV, using an all ions fragmentation approach with frames alternating between high and low fragmentation. All data were acquired using a scan range of *m/z* 50-1700 in positive and negative ionization.

**Lab C: LC-IMS-MS data analysis**

*Initial conversion.* Data was converted from vendor format to mzML for processing by DEIMoS (Data Processing for Integrated Multidimensional Spectrometry), an open-source Python package amenable to high dimensional mass spectrometry acquisitions, such as the liquid chromatography – ion mobility spectrometry – tandem mass spectrometry employed here.

*Peak detection.* An initial assessment of appropriate peak detection parameters was first performed by sampling the acquired three-dimensional (3D) space in each dimension (i.e. retention time, drift time, *m/z*) for representative features. The full width at half maximum was assessed per feature, per dimension to inform selection of peak detection parameters. Peak detection was then performed per data file at each mass spectrometry level (MS1 and MS2 at apparent collision energy) in the native acquisition dimension (i.e. 3D: retention time, drift time, *m/z*). Peak coordinates were returned as per-dimension coordinates and peak apex intensity. Each peak apex was interpolated to achieve sub-acquisition resolution to further refine coordinates in *m/z* and drift time dimensions. This operation was performed per feature detected, wherein a univariate spline of degree 2 was fit to each of the 1D projections of the 3D feature. The coordinate of the maximum interpolated intensity value was updated accordingly, per dimension.

*Collision cross section calibration*. Drift time values were converted to collision cross section (CCS) values using DEIMoS, which implements the Stow equation for CCS calibration. After linearization, a least-squares regression was performed to produce CCS versus drift time, from which the calibration parameters *beta* and *tfix* were derived. These parameters were then used to convert feature drift times to CCS per ionization mode. Note that the CCS calculation requires knowledge of nominal charge state; we assumed a nominal charge of 1 until refinement otherwise during the annotation phase of this procedure.

*Reference-based Alignment.* Internal standards were used to align retention times of data acquired on LC-IMS-MS and LC-IMS platforms. A support vector regression model was constructed for each LC-IMS-MS data file by putatively matching *m/z* values of detected internal standards against the LC-MS consensus list, per ionization mode. The fit transformation was then applied to all features detected on the LC-IMS-MS platform, per data file.

*Cross-sample Alignment*. Correspondence of features across samples was determined by an agglomerative clustering-based approach, as implemented in DEIMoS, with per-dimension tolerances informed by FWHM values discerned during peak detection parameter selection. We first aligned replicates (whether technical or analytical), keeping only those features that appear in all replicates to ensure reproducibility, aggregated by median feature coordinate. Remaining features were aligned across all samples, per ionization mode, and again aggregated by median feature coordinate. Aligned features were then matched to the LC-MS consensus list with tolerances of +- 10 ppm in *m/z* and +- 0.1 minutes in retention time.

**Alignment of Common Data Acquisition In-House Data Processed Features using MetabCombiner**

Each of the participating laboratories in this study provided individualized pre-processed feature tables, based on software and parameters described in Supplemental Methods. The features of these tables were aligned using the metabCombiner R package. As part of this workflow, one feature table (laboratory A) is used as a reference list and features from all other tables are sequentially aligned in a pairwise manner to the results of the previous alignment step. Feature filtering was kept to a minimum to retain as many features as possible for alignment: 50% presence in at least one specimen type, no features within {0.0025 Da, 0.05 minutes} within the same feature list. In one feature list (laboratory C), a retention time range filter was applied between 0.3 - 14 minutes for ease of RT mapping.

The table below lists the parameters associated with the main metabCombiner workflow steps: retention time ordered pair selection for RT mapping (selectAnchors), Generalized Additive Model fitting (fit_gam), pairwise similarity scoring (calcScores), and paired feature table reduction (reduceTable). A common m/z grouping value of 0.0075 Da was employed for initial construction. Parameter values not listed in this table may be assumed as the program defaults. After each pairwise feature list alignment, non-intersected entities are re-incorporated into the table (updateTables).

| | Ionization Mode | Positive | | | Negative | | |
|---|---|---|---|---|---|---|---|
| | Paired Laboratory | B | C | D | B | C | D |
| selectAnchors | tolmz | 0.003 | 0.002 | 0.002 | 0.003 | 0.002 | 0.003 |
| | tolQ | 0.3 | 0.3 | 0.3 | 0.3 | 0.3 | 0.3 |
| | tolrtq | 0.1 | 0.1 | 0.1 | 0.1 | 0.1 | 0.2 |
| | windx | 0.025 | 0.03 | 0.02 | 0.025 | 0.03 | 0.03 |
| | windy | 0.02 | 0.03 | 0.03 | 0.02 | 0.03 | 0.02 |
| fit_gam | k | 14 | 14 | 14 | 14 | 14 | 14 |
| | iterFilter | 2 | 2 | 1 | 2 | 2 | 1 |
| | rtx | (min, 11) | (min, 10) | (min, max) | (min, 9) | (min, 11) | (min, 9) |
| | rty | (min, 11) | (min, 11) | (min, max) | (min, 9.5) | (min, 12) | (min, 10) |
| calcScores | A | 70 | 70 | 70 | 70 | 70 | 70 |
| | B | 15 | 12 | 14 | 15 | 12 | 14 |
| | C | 0.2 | 0.2 | 0.1 | 0.2 | 0.2 | 0.1 |
| reduceTable | maxRTerr | 0.5 | 0.5 | 0.5 | 0.5 | 0.5 | 0.5 |
| | delta | 0.2 | 0.2 | 0.2 | 0.2 | 0.2 | 0.2 |

**Table S1:** 76 lipid internal standards from 18 classes (Avanti UltimateSPLASH™ ONE mixture and supplemental compounds)

| # | Internal Standard Name | Long Name | Class | Formula |
|---|---|---|---|---|
| 1 | 14:0-13:0-14:0 TG-d5 | 1,3-ditetradecanoyl-2-tridecanoyl-glycerol-d5 | TG | C44H79D5O6 |
| 2 | 14:0-15:1-14:0 TG-d5 | 1,3-ditetradecanoyl-2-pentadec-10-enoyl-glycerol-d5 | TG | C46H81D5O6 |
| 3 | 14:0-17:1-14:0 TG-d5 | 1,3-ditetradecanoyl-2-heptadec-10-enoyl-glycerol-d5 | TG | C48H85D5O6 |
| 4 | 16:0-15:1-16:0 TG-d5 | 1,3-dihexadecanoyl-2-pentadec-10Z-enoyl-glycerol-d5 | TG | C50H89D5O6 |
| 5 | 16:0-17:1-16:0 TG-d5 | 1,3-dihexadecanoyl-2-heptadec-10Z-enoyl-glycerol-d5 | TG | C52H93D5O6 |
| 6 | 16:0-19:2-16:0 TG-d5 | 1,3-dihexadecanoyl-2-nonadeca-10Z,13Z-dienoyl-glycerol-d5 | TG | C54H95D5O6 |
| 7 | 18:1-17:1-18:1 TG-d5 | 1,3-dioleoyl-2-heptadec-10Z-enoyl-glycerol-d5 | TG | C56H97D5O6 |
| 8 | 18:1-19:2-18:1 TG-d5 | 1,3-dioleoyl-2-nonadeca-10Z,13Z-dienoyl-glycerol-d5 | TG | C58H99D5O6 |
| 9 | 18:1-21:2-18:1 TG-d5 | 1,3-dioleoyl-2-heneicosa-12Z,15Z-dienoyl-glycerol-d5 | TG | C60H103D5O6 |
| 10 | 14:1 cholesteryl-d7 ester | cholesteryl-d7 myristoleate | CE | C41H63D7O2 |
| 11 | 16:1 cholesteryl-d7 ester | cholesteryl-d7 palmitoleate | CE | C43H67D7O2 |
| 12 | 18:1 cholesteryl-d7 ester | cholesteryl-d7 oleate | CE | C45H71D7O2 |
| 13 | 20:3 cholesteryl-d7 ester | cholesteryl-d7 eicosa-11,14,17-trienoate | CE | C47H71D7O2 |
| 14 | 22:4 cholesteryl-d7 ester | cholesteryl-d7 docosa-7,10,13,16-tetraenoate | CE | C49H73D7O2 |
| 15 | C16:1 Ceramide-d7 (d18:1-d7/16:1) | N-palmitoleoyl-D-erythro-sphingosine-d7 | Cer | C34H58D7NO3 |
| 16 | C18:1 Ceramide-d7 (d18:1-d7/18:1) | N-oleoyl-D-erythro-sphingosine-d7 | Cer | C36H62D7NO3 |
| 17 | C20:1 Ceramide-d7 (d18:1-d7/20:1) | N-eicos-11-enoyl-D-erythro-sphingosine-d7 | Cer | C38H66D7NO3 |
| 18 | C22:1 Ceramide-d7 (d18:1-d7/22:1) | N-erucoyl-D-erythro-sphingosine (d7) | Cer | C40H70D7NO3 |
| 19 | C24:1 Ceramide-d7 (d18:1-d7/24:1) | N-nervonoyl-D-erythro-sphingosine (d7) | Cer | C42H74D7NO3 |
| 20 | 16:1 SM (d18:1/16:1)-d9 | N-palmitoleoyl-D-erythro-sphingosylphosphorylcholine-d9 | SM | C39H68D9N2O6P |
| 21 | 18:1 SM (d18:1/18:1)-d9 | N-oleoyl-D-erythro-sphingosylphosphorylcholine-d9 | SM | C41H72D9N2O6P |
| 22 | 20:1 SM (d18:1/20:1)-d9 | N-eicos-11-enoyl-D-erythro-sphingosylphosphorylcholine-d9 | SM | C43H76D9N2O6P |
| 23 | 22:1 SM (d18:1/22:1)-d9 | N-erucoyl-D-erythro-sphingosylphosphorylcholine-d9 | SM | C45H80D9N2O6P |
| 24 | 24:1 SM (d18:1/24:1)-d9 | N-Nervonoyl-D-erythro-sphingosylphosphorylcholine-d9 | SM | C47H84D9N2O6P |
| 25 | 17:0-14:1 PC-d5 | 1-heptadecanoyl-2-myristoleoyl-sn-glycero(d5)-3-phosphocholine | PC | C39H71D5NO8P |
| 26 | 17:0-16:1 PC-d5 | 1-heptadecanoyl-2-palmitoleoyl-sn-glycero(d5)-3-phosphocholine | PC | C41H75D5NO8P |
| 27 | 17:0-18:1 PC-d5 | 1-heptadecanoyl-2-oleoyl-sn-glycero(d5)-3-phosphocholine | PC | C43H79D5NO8P |
| 28 | 17:0-20:3 PC-d5 | 1-heptadecanoyl-2-eicosatrienoyl-sn-glycero(d5)-3-phosphocholine | PC | C45H79D5NO8P |
| 29 | 17:0-22:4 PC-d5 | 1-heptadecanoyl-2-docosatetraenoyl-sn-glycero(d5)-3-phosphocholine | PC | C47H81D5NO8P |
| 30 | 17:0-14:1 PE-d5 | 1-heptadecanoyl-2-myristoleoyl-sn-glycero(d5)-3-phosphoethanolamine | PE | C36H65D5NO8P |
| 31 | 17:0-16:1 PE-d5 | 1-heptadecanoyl-2-palmitoleoyl-sn-glycero(d5)-3-phosphoethanolamine | PE | C38H69D5NO8P |

| | | | | |
|---|---|---|---|---|
| 32 | 17:0-18:1 PE-d5 | 1-heptadecanoyl-2-oleoyl-sn-glycero(d5)-3-phosphoethanolamine | PE | C40H73D5NO8P |
| 33 | 17:0-20:3 PE-d5 | 1-heptadecanoyl-2-eicosatrienoyl-sn-glycero(d5)-3-phosphoethanolamine | PE | C42H73D5NO8P |
| 34 | 17:0-22:4 PE-d5 | 1-heptadecanoyl-2-docosatetraenoyl-sn-glycero(d5)-3-phosphoethanolamine | PE | C44H75D5NO8P |
| 35 | 17:0-14:1 PG-d5 | 1-heptadecanoyl-2-myristoleoyl-sn-glycero(d5)-3-phospho-(1'-rac-glycerol)(sodium salt) | PG | C37H66D5O10P |
| 36 | 17:0-16:1 PG-d5 | 1-heptadecanoyl-2-palmitoleoyl-sn-glycero(d5)-3-phospho-(1'-rac-glycerol)(sodium salt) | PG | C39H70D5O10P |
| 37 | 17:0-18:1 PG-d5 | 1-heptadecanoyl-2-oleoyl-sn-glycero(d5)-3-phospho-(1'-rac-glycerol)(sodium salt) | PG | C41H74D5O10P |
| 38 | 17:0-20:3 PG-d5 | 1-heptadecanoyl-2-eicosatrienoyl-sn-glycero(d5)-3-phospho-(1'-rac-glycerol)(sodium salt) | PG | C43H74D5O10P |
| 39 | 17:0-22:4 PG-d5 | 1-heptadecanoyl-2-docosatetraenoyl-sn-glycero(d5)-3-phospho-(1'-rac-glycerol)(sodium salt) | PG | C45H76D5O10P |
| 40 | 17:0-14:1 PS-d5 | 1-heptadecanoyl-2-myristoleoyl-sn-glycero(d5)-3-phospho- L-serine (sodium salt) | PS | C37H65D5NO10P |
| 41 | 17:0-16:1 PS-d5 | 1-heptadecanoyl-2-palmitoleoyl-sn-glycero(d5)-3-phospho- L-serine (sodium salt) | PS | C39H69D5NO10P |
| 42 | 17:0-18:1 PS-d5 | 1-heptadecanoyl-2-oleoyl-sn-glycero(d5)-3-phospho- L-serine (sodium salt) | PS | C41H73D5NO10P |
| 43 | 17:0-20:3 PS-d5 | 1-heptadecanoyl-2-eicosatrienoyl-sn-glycero(d5)-3-phospho- L-serine (sodium salt) | PS | C43H73D5NO10P |
| 44 | 17:0-22:4 PS-d5 | 1-heptadecanoyl-2-docosatetraenoyl-sn-glycero(d5)-3-phospho- L-serine (sodium salt) | PS | C45H75D5NO10P |
| 45 | 17:0-14:1 DG-d5 | 1-heptadecanoyl-2-myristoleoyl-sn-glycerol(d5) | DG | C34H59D5O5 |
| 46 | 17:0-16:1 DG-d5 | 1-heptadecanoyl-2-palmitoleoyl-sn-glycerol(d5) | DG | C36H63D5O5 |
| 47 | 17:0-18:1 DG-d5 | 1-heptadecanoyl-2-oleoyl-sn-glycerol(d5) | DG | C38H67D5O5 |
| 48 | 17:0-20:3 DG-d5 | 1-heptadecanoyl-2-eicosatrienoyl-sn-glycerol(d5) | DG | C40H67D5O5 |
| 49 | 17:0-22:4 DG-d5 | 1-heptadecanoyl-2-docosatetraenoyl-sn-glycerol(d5) | DG | C42H69D5O5 |
| 50 | 17:0-14:1 PI-d5 | 1-heptadecanoyl-2-myristoleoyl-sn-glycero(d5)-3-phosphoinositol (ammonium salt) | PI | C40H70D5O13P |
| 51 | 17:0-16:1 PI-d5 | 1-heptadecanoyl-2-palmitoleoyl-sn-glycero(d5)-3-phosphoinositol (ammonium salt) | PI | C42H74D5O13P |
| 52 | 17:0-18:1 PI-d5 | 1-heptadecanoyl-2-oleoyl-sn-glycero(d5)-3-phosphoinositol (ammonium salt) | PI | C44H78D5O13P |
| 53 | 17:0-20:3 PI-d5 | 1-heptadecanoyl-2-eicosatrienoyl-sn-glycero(d5)-3-phosphoinositol (ammonium salt) | PI | C46H78D5O13P |
| 54 | 17:0-22:4 PI-d5 | 1-heptadecanoyl-2-docosatetraenoyl-sn-glycero(d5)-3-phosphoinositol (ammonium salt) | PI | C48H80D5O13P |
| 55 | 15:0 Lyso PI-d5 | 1-pentadecanoyl-2-hydroxy-sn-glycero(d5)-3-phosphoinositol (ammonium salt) | LPI | C24H42D5O12P |
| 56 | 17:0 Lyso PI-d5 | 1-heptadecanoyl-2-hydroxy-sn-glycero(d5)-3-phosphoinositol (ammonium salt) | LPI | C26H46D5O12P |
| 57 | 19:0 Lyso PI-d5 | 1-nonadecanoyl-2-hydroxy-sn-glycero(d5)-3-phosphoinositol (ammonium salt) | LPI | C28H50D5O12P |
| 58 | 15:0 Lyso PS-d5 | 1-pentadecanoyl-2-hydroxy-sn-glycero(d5)-3-phospho-L-serine (sodium salt) | LPS | C21H37D5NO9P |
| 59 | 17:0 Lyso PS-d5 | 1-heptadecanoyl-2-hydroxy-sn-glycero(d5)-3-phospho-L-serine (sodium salt) | LPS | C23H41D5NO9P |
| 60 | 19:0 Lyso PS-d5 | 1-nonadecanoyl-2-hydroxy-sn-glycero(d5)-3-phospho-L-serine (sodium salt) | LPS | C25H45D5NO9P |

| | | | | |
|---|---|---|---|---|
| 61 | 15:0 Lyso PG-d5 | 1-pentadecanoyl-2-hydroxy-sn-glycero(d5)-3-phospho-(1’-rac-glycerol)(sodium salt) | LPG | C21H38D5O9P |
| 62 | 17:0 Lyso PG-d5 | 1-heptadecanoyl-2-hydroxy-sn-glycero(d5)-3-phospho-(1’-rac-glycerol)(sodium salt) | LPG | C23H42D5O9P |
| 63 | 19:0 Lyso PG-d5 | 1-nonadecanoyl-2-hydroxy-sn-glycero(d5)-3-phospho-(1’-rac-glycerol)(sodium salt) | LPG | C25H46D5O9P |
| 64 | 15:0 Lyso PC-d5 | 1-pentadecanoyl-2-hydroxy-sn-glycero(d5)-3-phosphocholine | LPC | C23H43D5NO7P |
| 65 | 17:0 Lyso PC-d5 | 1-heptadecanoyl-2-hydroxy-sn-glycero(d5)-3-phosphocholine | LPC | C25H47D5NO7P |
| 66 | 19:0 Lyso PC-d5 | 1-nonadecanoyl-2-hydroxy-sn-glycero(d5)-3-phosphocholine | LPC | C27H51D5NO7P |
| 67 | 15:0 Lyso PE-d5 | 1-pentadecanoyl-2-hydroxy-sn-glycero(d5)-3-phosphoethanolamine | LPE | C20H37D5NO7P |
| 68 | 17:0 Lyso PE-d5 | 1-heptadecanoyl-2-hydroxy-sn-glycero(d5)-3-phosphoethanolamine | LPE | C22H41D5NO7P |
| 69 | 19:0 Lyso PE-d5 | 1-nonadecanoyl-2-hydroxy-sn-glycero(d5)-3-phosphoethanolamine | LPE | C24H45D5NO7P |
| 70 | 10:0 AC-d3 | [D3]decanoyl-L-carnitine | AC | C17H30D3NO4 |
| 71 | 12:0 AC-d3 | [D3]dodecanoyl-L-carnitine | AC | C19H34D3NO4 |
| 72 | 18:0 AC-d3 | [D3]octadecanoyl-L-carnitine | AC | C25H46D3NO4 |
| 73 | 16:0 FA-d3 | palmitic acid d3 | FA | C16H29D3O2 |
| 74 | 18:1 FA-d9 | oleic acid-d9 | FA | C18H25D9O2 |
| 75 | 20:4 FA-d11 | arachidonic acid-d11 | FA | C20H21D11O2 |
| 76 | chol-d7 | Cholesterol D7 | Chol | C27H39OD7 |

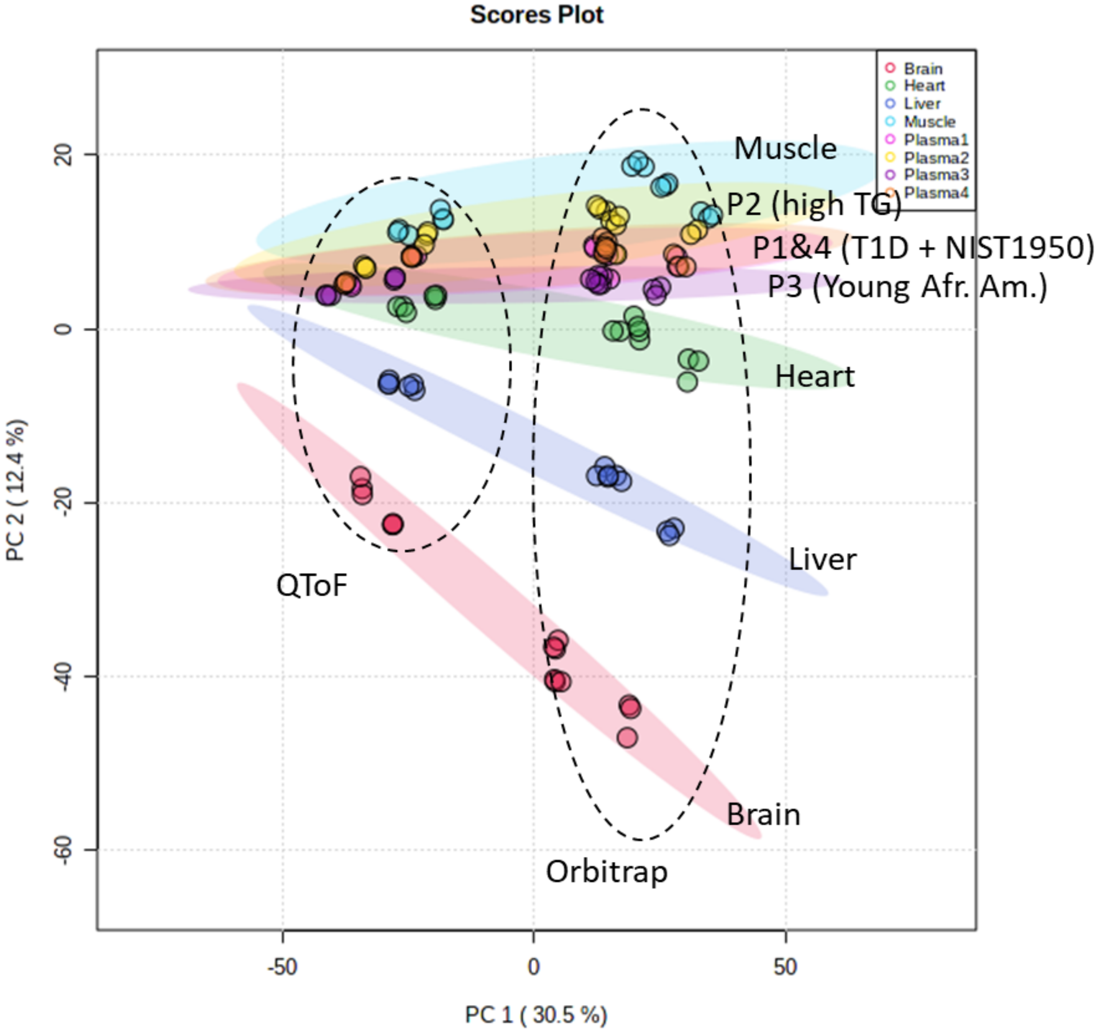


**Fig. S1**: Principal component analysis illustrating separation by sample type and by instrument type

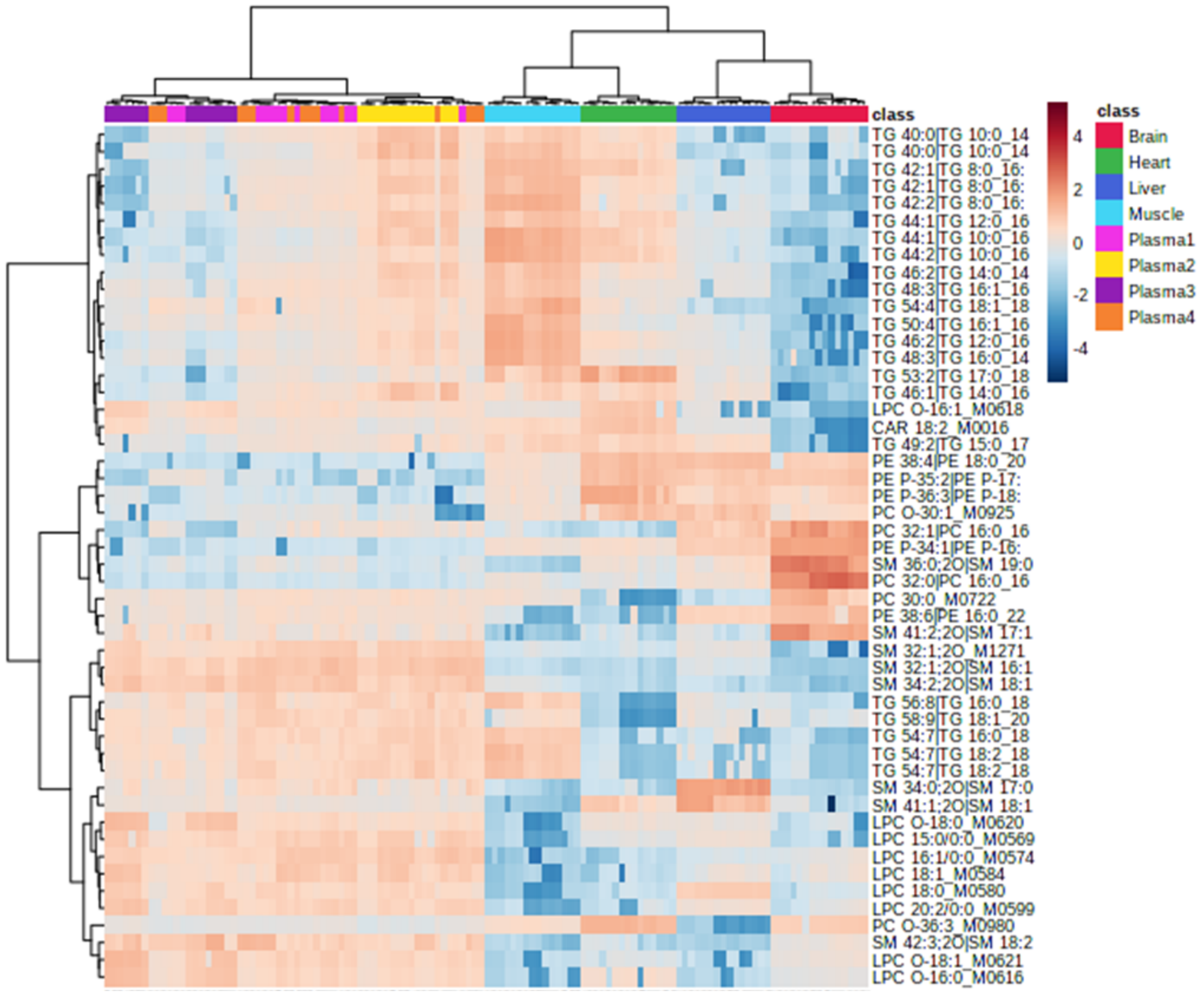


**Fig. S2**: Heatmap illustrating top differential identified lipids by sample type

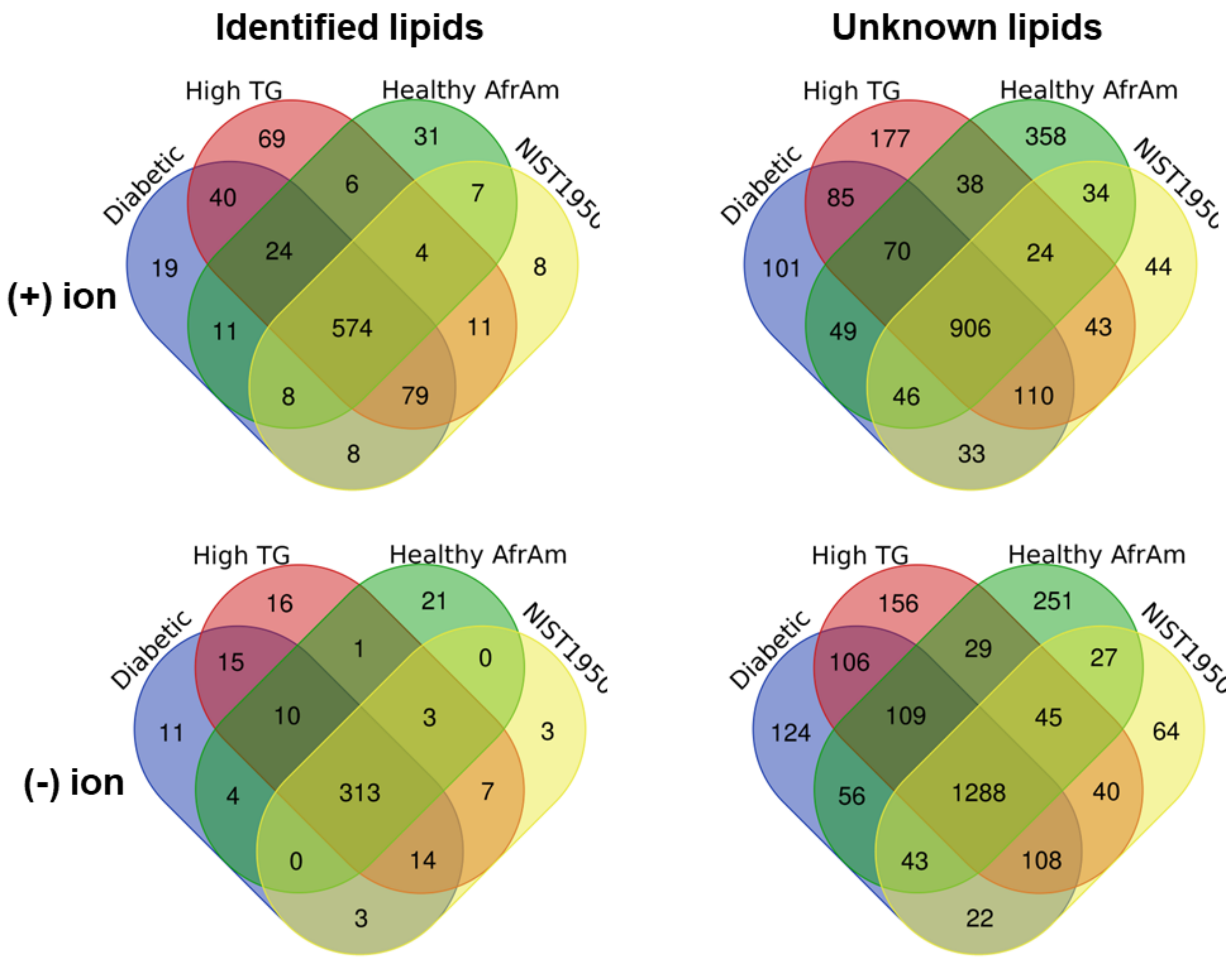


**Fig. S3: Overlap of identified and unknown lipids in plasma reference samples.** Data shown for features detected in >50% of common-method runs (including data from all study sites) of at least one sample type
